# *De novo* genome assembly of the olive fruit fly (*Bactrocera oleae*) developed through a combination of linked-reads and long-read technologies

**DOI:** 10.1101/505040

**Authors:** Haig Djambazian, Anthony Bayega, Konstantina T. Tsoumani, Efthimia Sagri, Maria-Eleni Gregoriou, Kristina Giorda, George Tsiamis, Kostas Bourtzis, Spyridon Oikonomopoulos, Ken Dewar, Deanna Church, Kostas D. Mathiopoulos, Jiannis Ragoussis

**Affiliations:** McGill University and Genome Quebec Innovation Centre, Department of Human Genetics, McGill University, Montreal, Canada; Department of Biochemistry and Biotechnology University of Thessaly Biopolis Larissa 41500, Greece; 10x Genomics 7068 Koll Center Parkway Suite 401 Pleasanton, CA 94566, USA; Department of Environmental and Natural Resources Management, University of Patras, Agrinio, Greece; Insect Pest Control Laboratory, Joint FAO/IAEA Division of Nuclear Techniques in Food and Agriculture, Vienna, Austria

**Keywords:** short-reads, linked-reads, and long-read technologies

## Abstract

Long-read sequencing has greatly contributed to the generation of high quality assemblies, albeit at a high cost. It is also not always clear how to combine sequencing platforms. We sequenced the genome of the olive fruit fly (*Bactrocera oleae*), the most important pest in the olive fruits agribusiness industry, using Illumina short-reads, mate-pairs, 10x Genomics linked-reads, Pacific Biosciences (PacBio), and Oxford Nanopore Technologies (ONT). The 10x linked-reads assembly gave the most contiguous assembly with an N50 of 2.16 Mb. Scaffolding the linked-reads assembly using long-reads from ONT gave a more contiguous assembly with scaffold N50 of 4.59 Mb. We also present the most extensive transcriptome datasets of the olive fly derived from different tissues and stages of development. Finally, we used the Chromosome Quotient method to identify Y-chromosome scaffolds and show that the long-reads based assembly generates very highly contiguous Y-chromosome assembly.

JR is a member of the MinION Access Program (MAP) and has received free-of-charge flow cells and sequencing kits from Oxford Nanopore Technologies for other projects. JR has had no other financial support from ONT.

AB has received re-imbursement for travel costs associated with attending Nanopore Community meeting 2018, a meeting organized my Oxford Nanopore Technologies.

## Introduction

The advent of affordable next generation sequencing technology and analysis methodologies has prompted major eukaryotic genome sequencing efforts, including the 5000 arthropod genomes initiative (i5k)^1,2^, a community effort to provide the genomic sequences of 5000 insect or related arthropod species. In this project, the onus lies on individual labs with a specific interest in these genomes to organize the sequencing, analysis, and curating of their genome. Insect genomes can be problematic to assemble as a result of high polymorphism, small physical sizes that might not allow sufficient quantities of DNA to be isolated from a single individual, and difficulty or inability to breed for genome homozygosity^3^. Therefore, it is critical to establish methodological approaches that will allow the *de novo* sequencing of insect genomes at high quality and low cost if the i5k target is to be achieved.

Among insects, the Tephritidae family contains some of the most important agricultural pests world-wide, such as the Mediterranean fruit fly (*Ceratitis capitata*), the oriental fruit fly (*Bactrocera dorsalis*), the Mexican fruit fly (*Anastrepha ludens*), the Australian Q-fly (*Bactrocera tryoni*) and the olive fruit fly (*Bactrocera oleae*). Olive flies are the major pest of wild and commercially cultivated olives trees^4–6^ causing an estimated annual damage of USD 800 million per year^4–6^. This is mainly because an olive tree that is not protected will practically have all its fruits infested. Consequently, the distribution of olive flies is closely associated with the expansion area of the olive trees; mostly the Mediterranean basin and the Middle East, California and South Africa^5,7^. Its economic importance in olive producing countries and several particularities of the olive fly’s biology render this insect an important subject of molecular studies. For example, being strictly monophagous, olive fly larvae restrict their diet to the olive sap, adapting their physiology to a single nutritional resource and relying on intestinal symbionts to supply essential dietary components that are not supplied by the olive. Being consummate specialists, olive fly larvae may restrict and at the same time specialize their defenses to the plant host, the olive fruit. Such adaptations inevitably should be reflected in its genome. For example, the small physical size^8^ may be related to the insect’s limited adaptability, and the gene content that reflects its particular needs. This fly is however, currently poorly characterized at a genomics level. Our previous work estimated the haploid genome size to be 322 Mb^8^. The genome also diploid and composed in 6 chromosomes with males being the heterogametic XY^8,9^.

*De novo* genome assembly which involves the reconstruction of the original contiguous DNA sequences from a collection of randomly sampled sequenced fragments (reads) is an important first step to fully understand the biology of an organism. However, this process is complicated by 3 main issues; presence of haplotype differences, repeated regions of the genome, and sequencing errors. The ideal sequencing platform should provide very long reads (in order of megabases) with single base-pair resolution, very low error rate, and low cost. However, no such platform currently exists. Short-read sequencing technologies such as ‘single nucleotide fluorescent base extension with reversible terminators^10^’ commonly referred to as Illumina sequencing (Illumina Inc) which deliver massive numbers of reads with very low error rates and at relatively low cost produce short reads of  ∼50-300 bp and *de novo* genome assemblies from such technologies are often so fragmented. On the other hand, long-read sequencing technologies such as protein nanopore DNA sequencing (nanopore sequencing, Oxford Nanopore Technologies) and Single Molecule Real-Time (SMRT) sequencing (PacBio sequencing, Pacific Biosciences) which deliver long reads have low throughput thus higher sequencing costs and have high raw-read error rates. However, assemblies from these technologies are much more contiguous yielding completely closed genome assemblies for small organisms like prokaryotes^11^.

The linked-reads technology^12,13^ from 10x Genomics leverages Illumina sequencing to produce pseudo-long reads by capturing a single ultra-long DNA fragment into an oil emulsion droplet and sampling along the length of the fragment using oligo-nucleotides bearing the same molecular barcode for each oil emulsion. Pooling and sequencing of all barcoded oligos and computationally linking all oligos taken from the same DNA molecule using the bespoke Supernova assembly tool ^14^provides a new powerful approach to using short-read technologies in *de novo* genome assembly and other applications in genomics. However, this entire methodology is optimized around human genomes and genomes of similar size, while for genomes of significantly smaller sizes, it is not clear whether the methodology will yield data superior to standard short read sequencing approaches.

Among organisms that employ an X-Y chromosome system, the Y chromosome has been notoriously difficult to assemble due to its heterochromatic and repetitive nature. For example, 80% of the *Drosophila melanogaster* Y chromosome is made up of repeats^15^. In most genome sequencing projects the Y chromosome sequence is fragmented into a large number of small, unmapped scaffolds^16^. Additionally, only a few genes reside on the Y chromosome and most of them are characterized by the presence of small exons, gigantic introns and very little conservation among species even of the same family. Therefore, Y chromosome assembly presents a unique challenge. In the olive fly, the Y chromosome encompasses the male determining factor, M, the ‘Holy Grail’ of Tephritidae genetics for over 30 years ^17^. The M factor is the initial switch of the sex-determining cascade in tephritids, a switch that has been speculated to differ from the one used by the model dipteran Drosophila (for a review see ^18^). Unraveling the sex-determination cascade will shed light on the evolution of a major developmental pathway in most animals, as well as the evolution of the sex chromosomes themselves. Finally, the M factor can serve as a genetic sex-switching tool in environmentally friendly approaches of fly control that involve the release of sterilized males in the environment (Sterile Insect Technique, SIT)^19,20^. The olive fly Y chromosome is however, particularly small^21,22^, karyotypically appearing as a ∼4 Mb dot chromosome of *D. melanogaster*^23^.

To benefit from the pros of each sequencing technology, hybrid approaches that aim to sequence organisms using different approaches and then combine the data either at the level of error correction of long-reads and/or scaffolding and gap-closing of short-read based assemblies are increasingly widely applied (reviewed elsewhere^24^). Hybrid genome assemblies have shown more accuracy and contiguity^11,25^, and are now a preferred approach to *de novo* genome assembly. Here, we introduce the sequence of the whole genome of the olive fly, as part of the i5k initiative. We mainly focus on our technological approaches and particularly on the use of the linked-reads technology^13^ (10x Genomics, USA). We show a combination of different technologies: Illumina short reads and mate-pairs, and long reads from Pacific Biosciences (PacBio) and Oxford Nanopore Technologies (ONT). We also demonstrate that the 10x genomics assemblies can be combined with long reads derived using PacBio or Oxford Nanopore as well as mate pair libraries to produce significantly improved assemblies with a maximum NG50 length of 6.82Mb and an LG50 count of 15 scaffolds when all technologies are combined. Insights into the biology of the organism that stem from the analysis of its genome and various transcriptomes will be presented in future publications. We further aimed to identify chromosome Y-specific scaffolds and offer a significant hope for a more complete assembly of the Y chromosome that could eventually lead to the discovery of the elusive M factor.

## RESULTS

### Initial assembly using Illumina paired end, mate pair and PacBio reads

Our initial assembly was performed combining a series of libraries and sequencing technologies. First, a short-read assembly was created using two Illumina short insert libraries made separately from male and female flies (see Supplementary Table 1). Sequencing of the male DNA yielded 114.8 million 100bp paired-end reads resulting in11.4 Gb and 36X theoretical coverage of *B. oleae*. The respective number of reads, total bases and coverage for the female DNA sequencing were 197 million, 19.7 Gb, 61X. The male and female reads were combined and assembled together using a short read assembler Ray^26^using a kmer of k41 which produced the largest contig. This first step produced an assembly with a NG50 length of 13,362 in 5,068 contigs. Both male and female reads were also assembled independently but produced less good assemblies than mixed-reads assembly (results not shown). The final short read assembly was then scaffolded using the three mate pair libraries (see Supplementary Table S1) using SSPACE^27^. These mate-pair settings were then used with the reads to produce a scaffold with a NG50 length of 309,290 in 205 scaffolds.

**Table 1:**
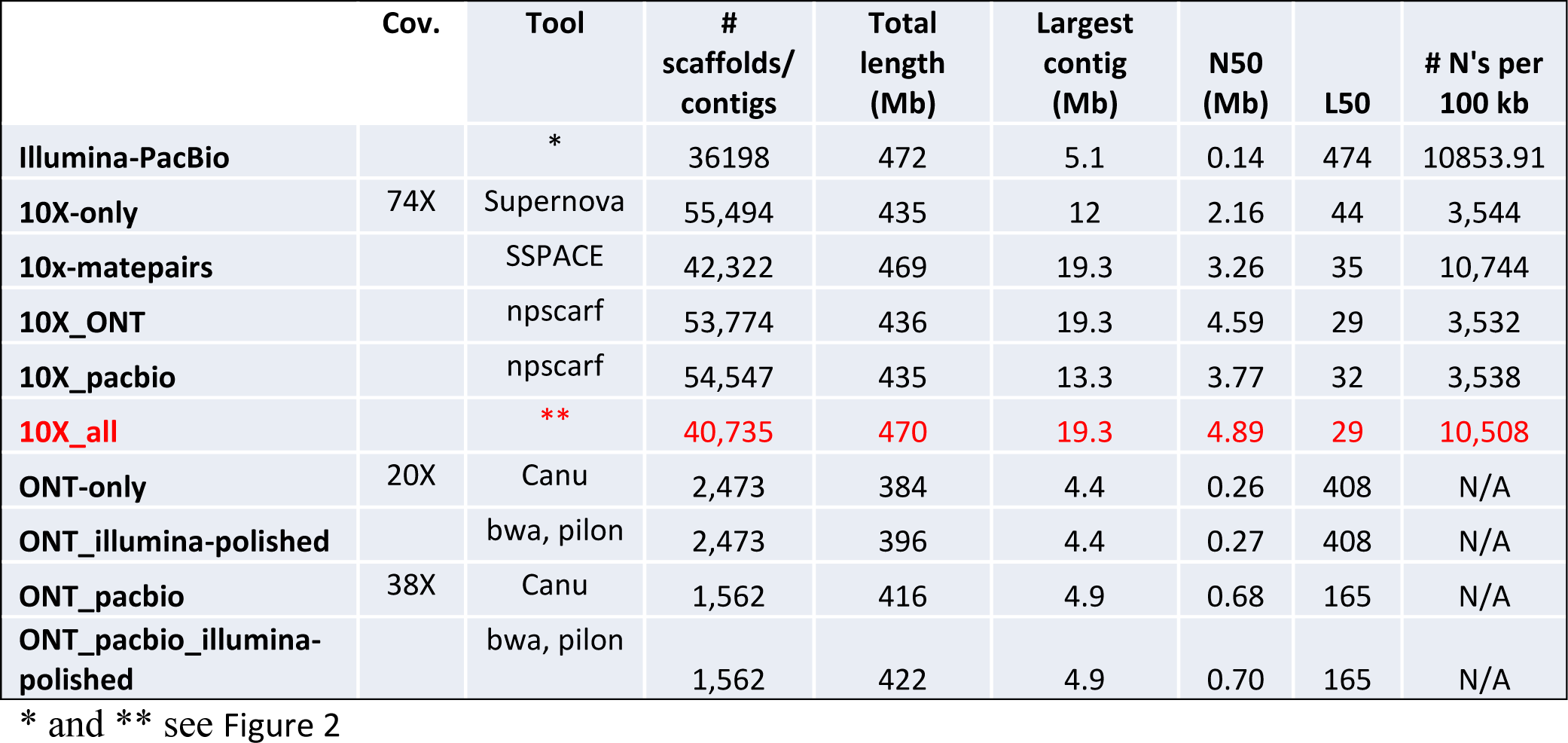
Assembly statistics for the 10 different assemblies generated. Quality metrics were generated using Quast^28^. Highlighted is red is the most contiguous assembly generated using linked reads assembly followed by gap-closing and scaffolding with illumina mate-pair reads, and long reads from PacBio and ONT.

We then performed PacBio Sequencing using high molecular weight DNA from male *B. oleae* which generated a 20X coverage using RS II technology. As a final step, PacBio reads were used to fill the sequence gaps left behind by the scaffolding process resulting in a decrease in gap bases from 44,015,646 bases to 27,804,509. This assembly was filtered for scaffolds with more than 10X of average Illumina coverage and a minimum length of 500bp and submitted to NCBI (GenBank assembly accession: GCA_001188975.2). The submitted assembly had a total length of 471,780,370 bases with a scaffold N50 length of 139,566bp reached with 474 scaffolds (Figure 1, Table S2). This assembly was submitted to i5k (https://i5k.nal.usda.gov/Bactrocera_oleae). This submission represented an improvement from the previous *Bactrocera oleae* assembly (submission GCA_001014625.1) which had a scaffold N50 length of 10,503bp reached with 6,242 scaffolds (see Supplementary Figure 1 A and B). This assembly is hereafter referred to as Illumina-PacBio. Table1 provides a summary of the quality features of all the assemblies.

**Figure 1:**
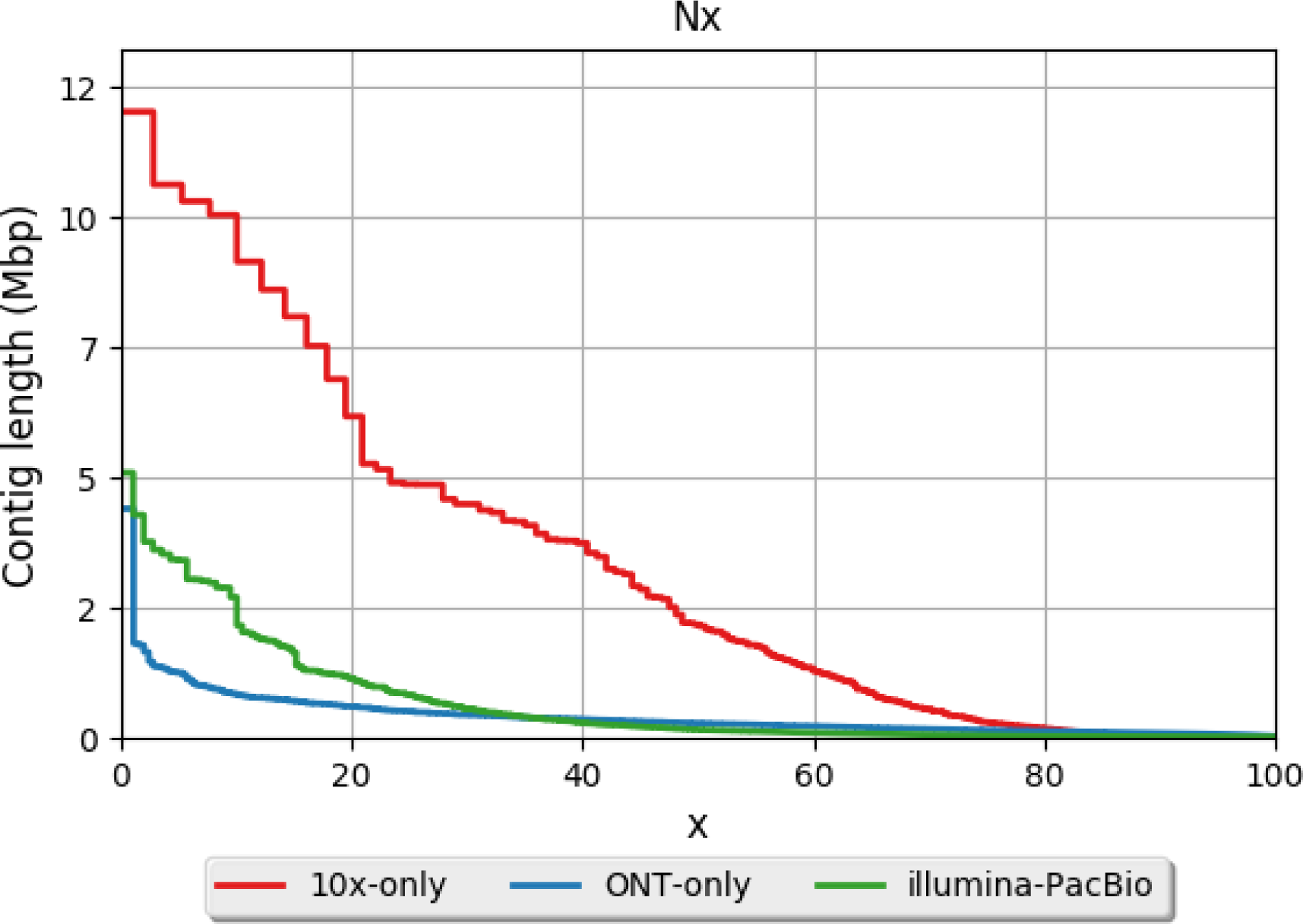
Contiguity plot generated using Quast^28^. Three assemblies were assessed; our initial assembly generated from Illumina short-read assembly followed by PacBio gap-closing (GenBank assembly accession: GCA_001188975.2, illumine-PacBio), an assembly generated only from Oxford Nanopore technology (ONT, ONT-only), and an assembly generated from the 10x Genomics linked-reads technology (10x-only). When all contigs from an assembly are ordered from largest to smallest, x is described as the value at which x% of the genome is contained in contigs at and above this contig length. The y-axis shows the contig/scaffold length at each of the Nx values. The N50 is the contig/scaffold length at which 50% of the genome is contained in scaffolds bigger as big as that contig/scaffold.

### Utilization of linked reads to generate a *Bactrocera oleae* assembly

The 10xGenomics platform which generates linked-reads has great potential to yield high quality assemblies both in terms of base accuracy and contiguity. The current configuration of the 10x Chromium system (DNA input amount, number of partitions, and sequencing depth) are optimized for the human genome and need to be modified for smaller genomes^14^. The key features of our sample preparation were the extraction of high molecular weight DNA, and size selection for DNA molecules >40kb reads (see materials and methods). A raw total of 414,094,826 reads were generated from which 22,793,359 barcodes were identified. However, 13,528,431 barcodes were rejected as low quality (see materials and methods) leaving 9,264,928. We then corrected the barcode sequences reducing the number of partitions to 3,700,604 containing 395,344,827 reads. The partitions were further filtered to retain only those that had >10 reads, leaving 1,443,720 barcodes containing 389,830,635 reads. Each read was then assigned to its partition. The Supernova assembler was then used to develop the *de novo* assembly following rounds of optimization by changing the number of partitions and coverage required to give the most contiguous assembly (as measured by the assembly NG50). The resulting genome was analysed using Quast^28^ (Figure 1, Table S2). The total assembly length is 434.81 Mb with a scaffold N50 of 2.15 Mb with the largest scaffold stretching 12 Mb. The L50 (representing the number of the largest scaffolds comprising 50% of the total assembly length) is only 44. This assembly is hereafter referred to as 10x-only. The 10x-only is also the most contiguous single (as opposed to hybrid) assembly we assembled. This assembly however, had 3543.94 gaps (N’s) per 100 kb.

### *De novo* assembly of the olive fly using Oxford Nanopore Technologies (ONT)

Because ONT current generates the longest raw reads of any commercially available sequencer with no theoretical limits^29^ there is potential to greatly increase assembly contiguity because the long reads can anchor repeated regions of the genome to unique adjacent regions making assemblies more contiguous. These features make the MinION very attractive for *de novo* genome assembly. We therefore, sought to utilize this technology to sequence the olive fly. High molecular weight DNA was extracted from a pool of adult male flies and used to prepare ONT sequencing libraries. Overall, 9 flow cells were used to perform the sequencing using 2 sequencing pore versions; R7.3 and R9.4. Over 95% of all the sequencing data was generated from the R9.4 nanopore chemistry which, together with improvements in sequencing software and basecalling algorithms, demonstrated much higher throughput. In total, the number of reads generated were 1,821,467 which yielded 9,517,723,896 bases. At the estimated *B. oleae* genome size of 322Mb, this represented a theoretical coverage of 28X. The read N50 was 11 kb with the longest read generated being780 kb.

Canu assembler^30^ was used to perform *de novo* genome assembly using ONT reads (ONT-only assembly) with “genome Size=320m” parameter. The resulting assembly was analysed by both custom scripts and Quast^28^. The final assembly contained 2,473 contigs with a total size of 384Mb. The assembly contig (not scaffold) N50 was 258 kb and the assembly L50 was 408 reads. The largest contig was 4.38Mb, and the assembly was generated without any ‘N’s and thus reflects true contigs and not scaffolds (Figure 1, Table S2).

We then attempted to generate hybrid assemblies and assess their features. Figure 2 shows a summary of the combinations tested. First, we combined the previously obtained 20X *B. oleae* coverage from PacBio sequencing and the 28X coverage of ONT sequencing and assembled it together using Canu. This long-read hybrid assembly (ONT-PacBio) yielded a bigger and more contiguous assembly with 1,562 contigs totaling 415.87Mb assembly size, N50 of 685.8 kb, and L50 of just 165 contigs (Figure 3). We also performed error correction of the ONT-only and ONT-PacBio assemblies by aligning Illumina short-reads onto these assemblies with BWA-MEM^31^ and processing the alignment files with Pilon^32^. These long-read assemblies were then aligned to the 10x-only assembly using MUMmer^33^ and alignment identity determined using the dnadiff tool of MUMmer. Indeed, alignment identities between the 10x-only assembly and ONT-only and ONT-PacBio assemblies improved from 96.49% to 98.99% and 97.92 to 99.26, respectively. The polishing of the genomes has only minimal improvements to other metrics analyzed (Table1).

**Figure 2:**
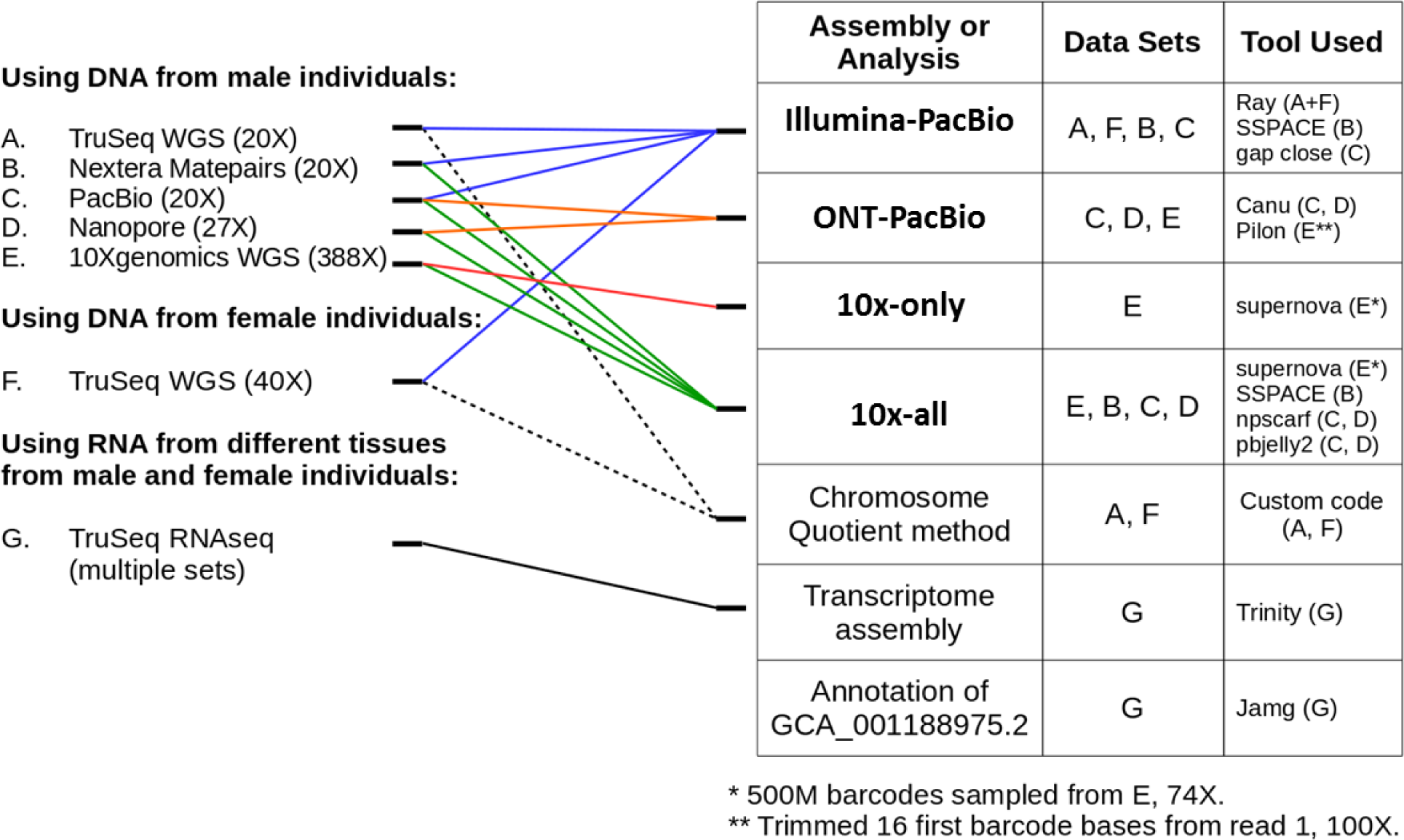
Schematic summary of the genome assembly tools used, datasets used, and the integration process.

**Figure 3:**
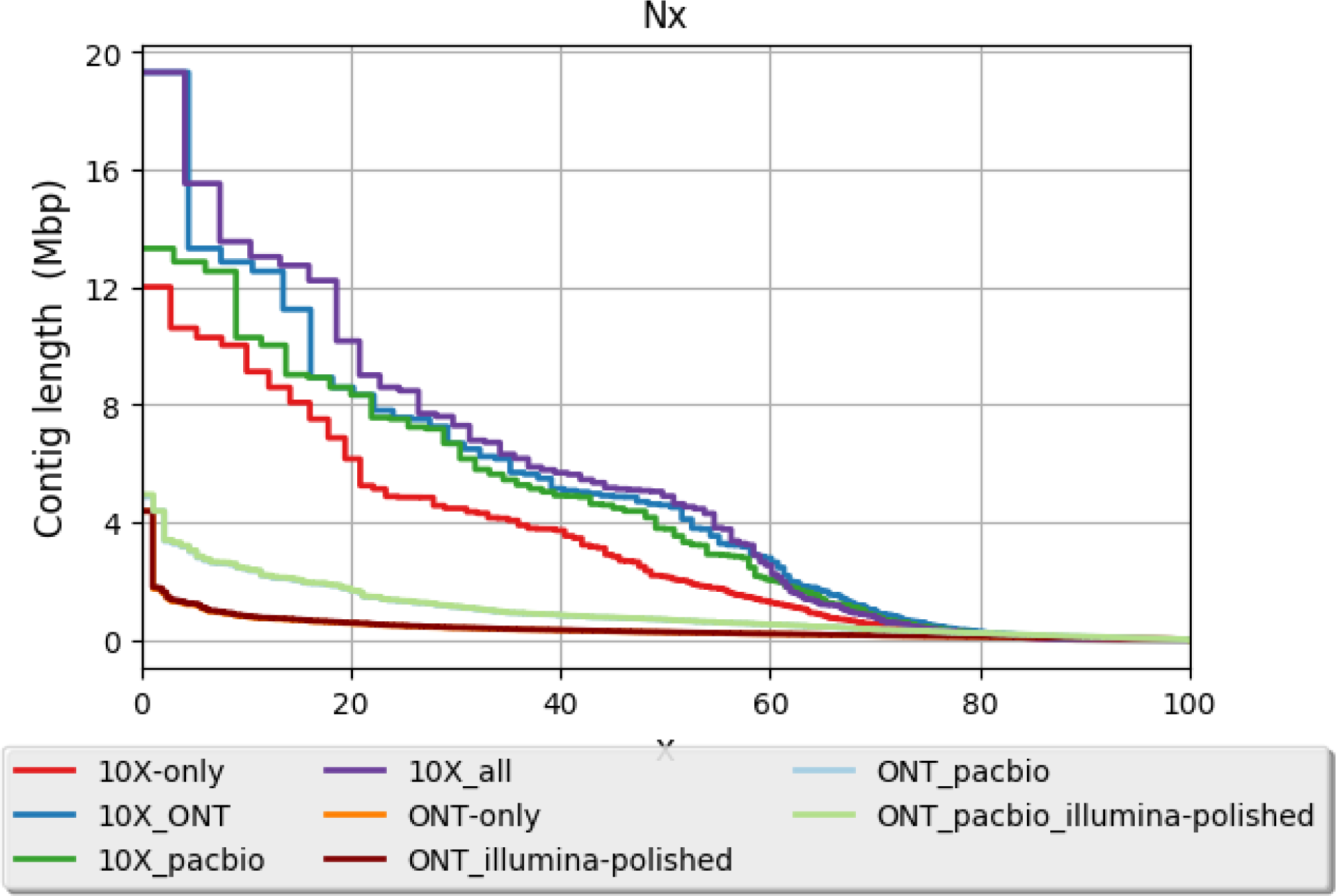
Contiguity plot comparing different assemblies. When all contigs from an assembly are ordered from largest to smallest, x is described as the value at which x% of the genome is contained in contigs at and above this contig length. The y-axis shows the contig/scaffold length at each of the Nx values. The N50 is the contig/scaffold length at which 50% of the genome is contained in scaffolds bigger as big as that contig/scaffold.

### Integration of linked-reads assembly with long-reads assembly

We attempted to increase the contiguity of the10x-only assembly further by scaffolding of its scaffolds and filling gaps using data from long-read sequencing technologies such as PacBio and Oxford Nanopore, as well mate-pair libraries (Figure 2). Using SSPACE, mate-pair sequences were used to scaffold the 10x-only assembly. This had a noticeable improvement on the 10x-only assembly increasing the N50 to 3.26 (51%), increasing the length of the longest scaffold from 12 Mb to 19.3 Mb. Mate-pair scaffolding however, increased the number of Ns in the assembly from 3543.94 to 10744.3 per 100 kb (Figure 3).

Scaffolding the 10x-only assembly using PacBio reads (20X coverage) using PBJelly increased the 10x-only scaffold N50 to 3.77 Mb (74% increase) and reduced the L50 to 2 scaffolds (Figure 3). Interestingly, scaffolding the 10x-only assembly using ONT reads (28X coverage) had the biggest improvement on contiguity. The scaffold N50 was more than doubled to 4.59 Mb (112% increment) and L50 reduced to 29 scaffolds (Figure 3). Further, the largest scaffold was increased to 19.3 Mb up from 12 Mb. Interestingly, combination of all technologies by scaffolding the 10x-only assembly first with mate-pairs then PacBio followed by ONT had slight improvements to the assembly compred when compared with 10x-ONT assembly. The scaffold N50 was minimally increased to 4.89 Mb (6.5% increase over th 10x-ONT assembly), with no apparent effect on the L50 or size of the largest scaffold. This suggested that the combining *de novo* assemblies from 10x Genomics linked-reads with long-reads from Oxford Nanopore provides about as contiguous assemblies as would be obtained from combining linked reads, mate-pairs, and PacBio. We observed that the best aproach to combine datasets was determined to be a scaffolding approach followed by gap filling.

### Repeat content analysis

A survey of repeats in the 10x-only assembly was carried using 2 different methods; RepeatMasker and RepeatScout^34^ which yielded largely similar results (Figure 4). Whereas RepeatMasker uses a database of curated repeat elements, RepeatScout uses a consensus driven seed expansion algorithm that first finds a short seed repeated in the genome which is extended left and right until most of the alignments at the repeat locations stop aligning to the seed. This allows the method to catalog both full length repeats as well as the partial instances. RepeatMasker masked 11.96% of the genome marking it as constitutive of repetitive sequence. Both methods showed that transposons account for most repeats followed by unclassified elements, long interspersed nuclear elements (LINEs), long terminal repeat elements (LTR), and short interspersed nuclear elements (SINEs) (Figure 4).

**Figure 4:**
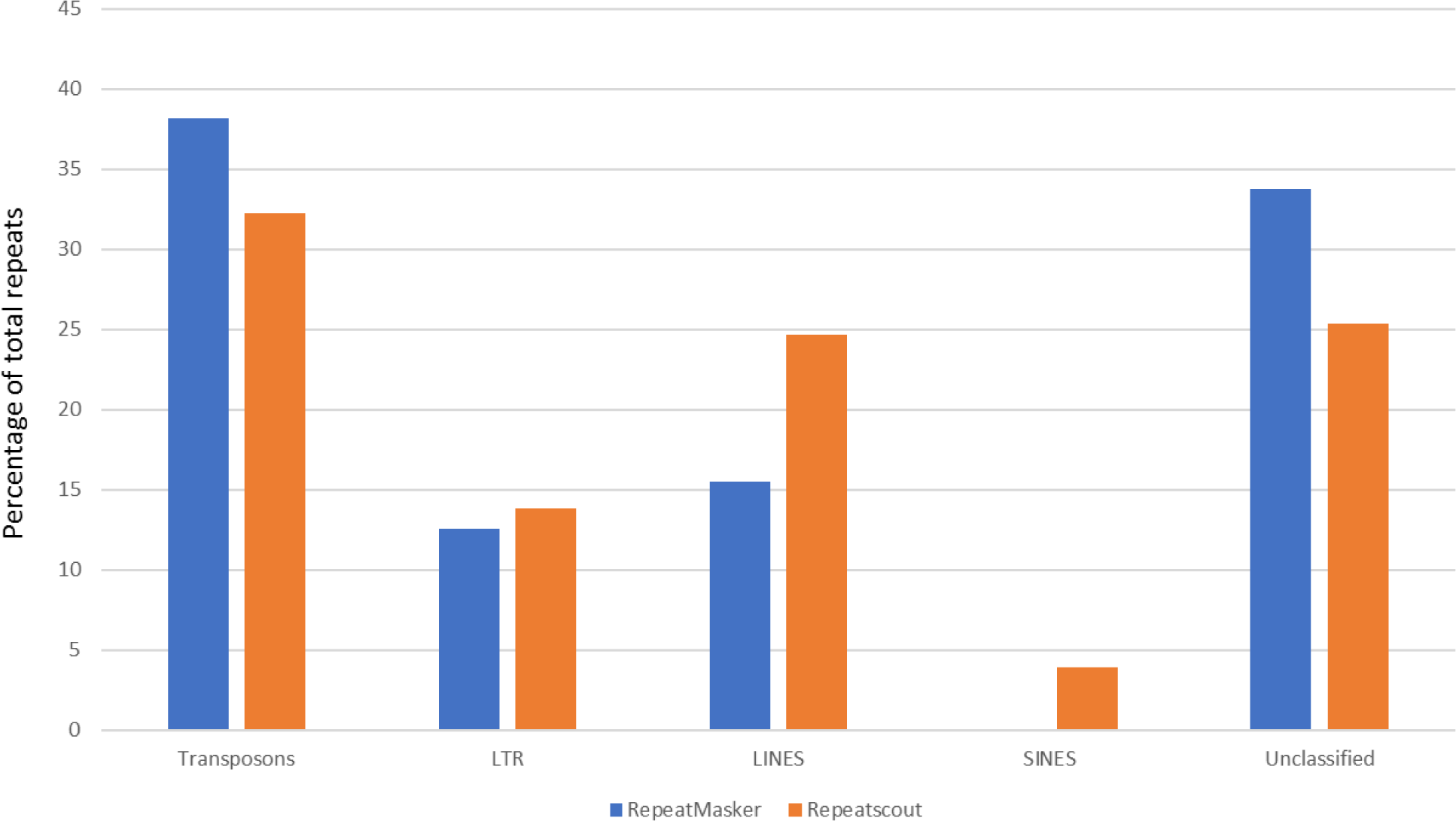
Distribution of different classes of repeat sequences in the olive fly genome. RepeatMasker was used to identify and mask repeat sequences using insect repeat database (Blue). Repetitive sequences were also identified using Repeatscout^34^ which performs *de novo* repeat sequence identification (Orange). Repeat sequences identified were annotated using TEclass^50^.

A total of 8520 *de novo* repeats were found with RepeatScout in the 10x-only assembly. These ranged in length from 50bp to 8951bp and covering 34.48% or 150Mb out of the total assembled genome of 435 Mb (Supplementary Figure S1). On average, repeats occur 106 times in the genome with 12% of repeats occurring more than 100 times. Using the location of these repeats a hard-masked version of the assembly was produced for downstream Y chromosome scaffold detection. The 10x-only assembly captured majority of the repeats as shown by the similarity in number and size of repeats present in the linked read assembly and linked-reads assembly gap-closed with long-reads from PacBio (Supplementary Figure S1).

### *De novo* transcriptome of the olive fly using different tissues

RNA sequencing was performed from 15 tissues and/or organs; 7 from female, 3 from male and 5 of mixed origin. The tissues and/or organs included eggs, larvae, pupae, heads, testes among others (Supplementary Table3). Between 29 and 55 million reads per library were used to perform *de novo* transcript assembly using Trinity^35^. This produced 133,003 transcripts with a median transcript length of 503 bp (Supplementary Table 4 and Supplementary Figure S2). The completeness of the assembly was evaluated using gVolante^36^ which determines the recovery of Athropoda BUSCOs in the assembly. Of the 1066 BUSOCs searched 98.87% (1054) were recovered in the assembly (Supplementary Table S5 and Supplementary Figure S3) suggesting that the transcriptome captured the majority of genes.

### Evaluation of assembly completeness using transcriptome derived data

In order to evaluate the genome assemblies, we aligned the assembled transcripts to the genome assembly and evaluated the number of different transcripts that aligned to the assemblies, the splice site sequences of putative exons divided according to splice site sequence (including canonical and non canonical splice sites), as well as the number of such splice sites per assembly. The results show how well an independently assembled transcriptome from short-read data aligns back to our assemblies. The combined assembly of 10x-all recovered the highest number of transcripts, as well as the broadest spectrum of splice site sequences. The number of transcripts mapped as full length was systematically higher in the combined diploid linked-read assembly. We noticed that our original Illumina-PacBio assembly performs less well compared to linked-reads based assemblies.

We further evaluated the completeness of the assembly using a list of orthologous ancestral genes termedBasic Universal Single Copy Orthologs (BUSCOs)^37^. Indeed, across the 4 lineages analysed; eukaryota, arthropoda, insecta, and diptera, 99%, 99.2%, 98.8%, and 97.2% of the genes surveyed were captured in the 10x-ony assembly, compared to 6.9%, 12.9%, 16.3%, and 10.3% respectively, captured in the ONT-only assembly (Figure 5). However, error-correction of the ONT-only assembly using Illumina short-reads increased the number of genes captured to 95.7%, 96.4%, 95.7%, and 94.1%, for the eukaryota, arthropoda, insecta,and dipteralineages, respectively (Figure 5).

**Figure 5:**
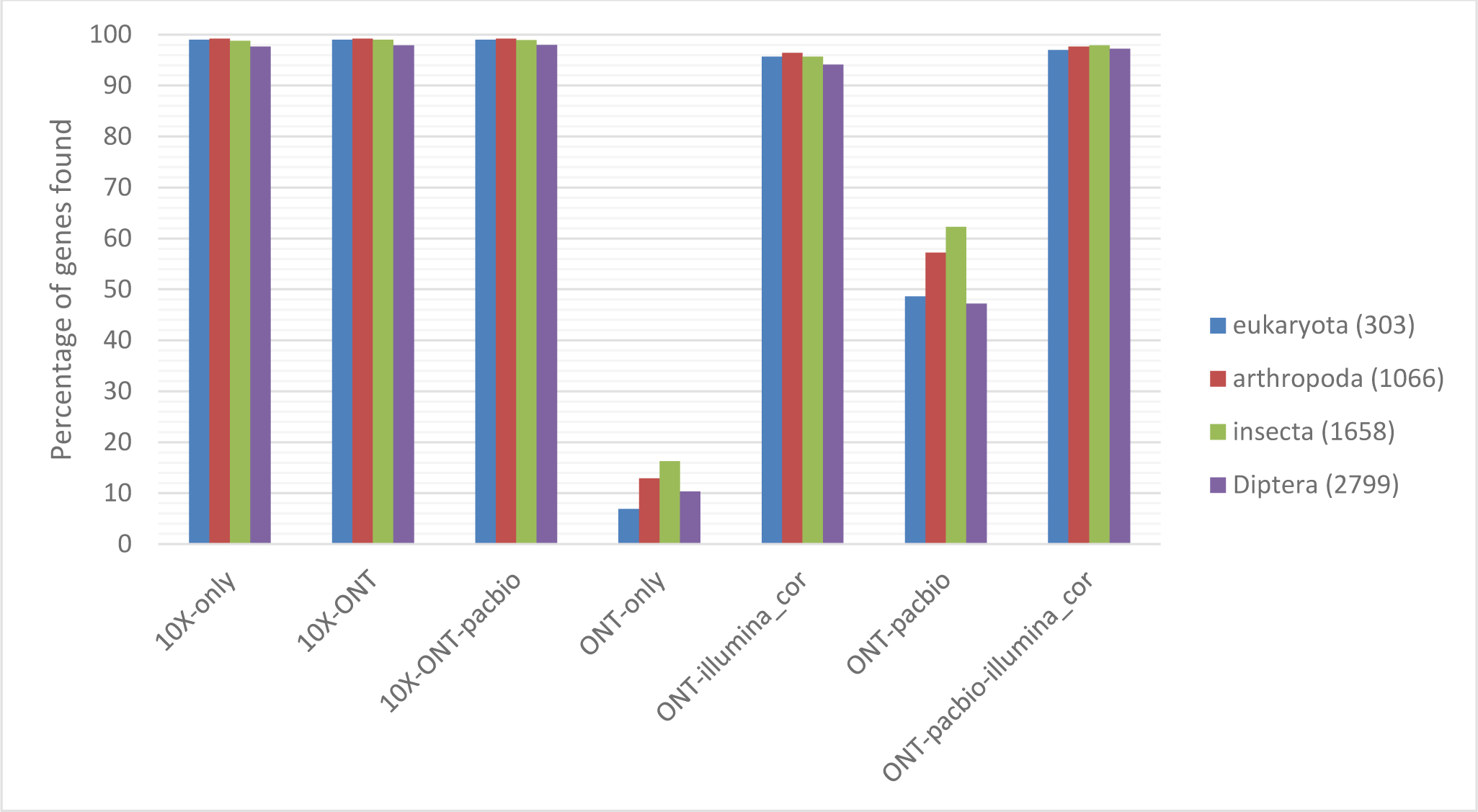
Bar graph of Basic Universal Single Copy Orthologs (BUSCOs)^37^ identified in the different assemblies. BUSCOs from Eukaryota (303), Arthropoda (1066), Insecta (1658) and Diptera (2799) were surveyed. ONT-illumina_cor and ONT-pacbio-illumina_cor are Illumina polished versions of ONT-only and ONT-PacBio assemblies.

### Identification of symbiont derived sequences in the assemblies

Sequences that belong to bacterial contaminants or symbionts were identified using an approach similar to the one applied to the Mediterraneanfruit fly^38^, using the Illumina-PacBio assembly (GCA_001188975.2). We identified small fragments that displayed homology with *Wolbachia* sequences. The biggest fragment identified was 855 bp in length. In total 7 fragments have been identified with a size range from 855-259 bp. No *Cardinium* and *Spiro* sequences were present either in the raw dataset (sra files) or in the assembled contigs.

Finally, our second approach using bacterial complete and draft genomes deposited in NCBI revealed the presence of a significant number of bacterial sequences that were filtered in the assembled scaffolds of the olive fly. The size of these sequences varied from 1103-100 bases. Alignments smaller than 100 bases were considered as noise and were not included in the analysis. The percentage of sequence identity was between 89%-70%. At least 109 sequences belonging to respective OTU/bacterial species were identified in the assembly scaffolds of *B. oleae*.

Using these results, we then downloaded the whole reference for all the observed hits and aligned these references to the scaffolds from the 10x-only assembly. We noticed that the corresponding scaffolds were small (less than 2Kb). As a result, these scaffold scan be discounted from the assembly. Setting a coverage cutoff of 90% (scaffold with more than 90% symbiont sequence) we identified two scaffolds that can be removed from the assembly. We also observe that for other larger scaffolds only a fraction contains sequence from other species and point to putative integrations from symbionts, therefore these scaffolds will be retained in the assembly.

We also searched for DNA contamination in the 10x-only reads and assembly using Kraken^39^. The 10x-only reads and scaffolds were searched against a database of human, virus, plasmids, archeal, fungi, and protozoa. About 1.35 % of reads and 7.7 % of scaffolds were determined to be contaminated (Supplementary Figure S4). Human DNA was the highest contaminant accounting for 62% of scaffold contaminants followed by Bacteria at 36%. Interestingly, the contamination was largely restricted to short scaffolds; for example, 84 % of all scaffolds showing contamination were less than 10 kb.

### Identification of sex chromosome sequences

In order to find putative X or Y chromosome scaffolds we used the Chromosome Quotient (CQ) method^40^. The CQ reflects the median ratio of female to male reads coverage when these reads are aligned to a male genome assembly. The CQ values will cluster around zero, one, and two for Y, autosome, and X scaffolds, respectively. Using the repeat masked versions of 3 genomes; the 10x-only, 10x-all, and ONT-PacBio, which were all generated from male olive fly DNA, male and female short Illumina reads (40X coverage of each) were independently mapped. For each dataset, we calculated the depth of coverage at each base for all scaffolds and filtered out positions with less than 10X of male coverage ensuring a minimum of evidence from male DNA. Then for each scaffold we calculated the CQ value for all positions. Considering only the scaffolds with a CQ of 0 we obtained a total length of putative Y of 3,827,631 bases with 1686 scaffolds from the 10x-only assembly. Increasing the CQ to the maximum allowed of 0.3, the length of scaffolds captured increased to 4,310,741. Although the latter setting of 0.3 gave us more putative Y scaffolds we performed a sanity check analysis where the male and female read sets were swapped to find positive “female Y” scaffolds (male to female ratio < 0.3). Here we found that using a CQ of 0.3 returned as much as 900kb of scaffolds whereas a cutoff of zero returned just 25kb. For this reason, the result from a cutoff of zero was conserved for downstream analysis. The Y chromosome scaffolds were evaluated using Quast (Supplementary Figure S5, Supplementary Table S6).

The 10x-all Y scaffolds showed the highest contiguity have a scaffold N50 of 60 kb and the largest scaffold being 318 kb (Figure 6, Supplementary Figure S5). The ONT-PacBio scaffolds also showed very high contiguity. Despite being the largest (4.72 Mb) the total number of contigs was only 125 compared to 873 scaffolds for a 3.9 Mb 10x-all Y chromosome assembly and 2138 scaffolds for a 3.8 Mb 10x-only Y chromosome assembly. The ONT-PacBio Y chromosome scaffolds do not contain gaps unlike the 10x-only and 10x-all assemblies which have 55238.99 and 75614.73 N’s per 100 kb, thus representing true contigs and not scaffolds. Capturing the 4.72 Mb Y chromosome in just 125 gapless contigs using only the long reads demonstrated the utility of long reads in assembling repeated regions like the Y chromosome which is known to be highly repeated.

**Figure 6:**
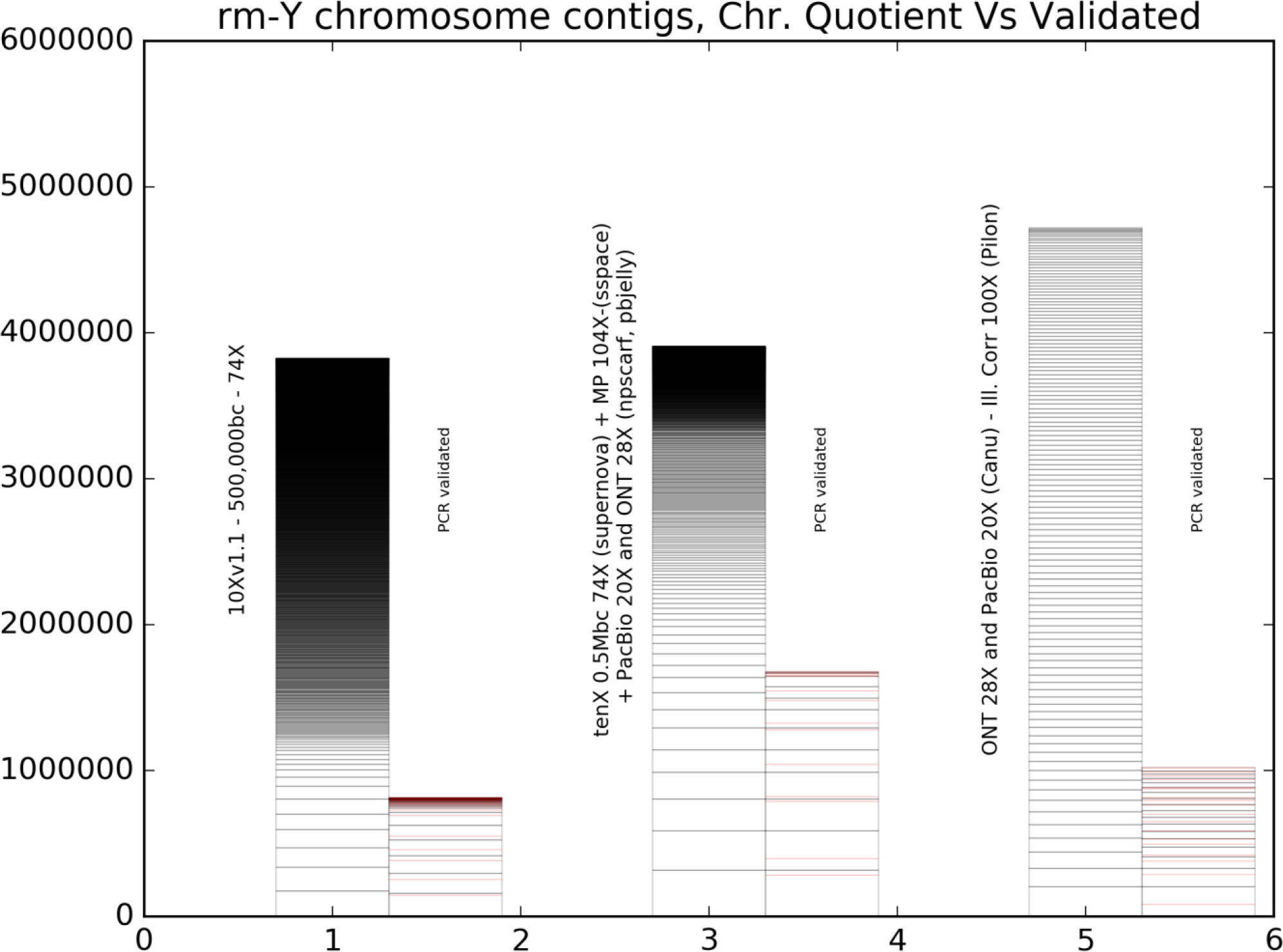
Plot showing Y chromosome scaffolds/contigs identified in 3 different assemblies using used the Chromosome Quotient (CQ) method^40^. The scaffolds/contigs are ordered from longest to shortest. For each assembly the total scaffolds/contigs are shown in black bars while the PCR validated scaffolds/contigs are shown in pink.

### Validation of Y-chromosome sequences by PCR

In order to validate that the Y scaffolds identified using the CQ method were indeed Y-linked scaffolds, 85 primer pairs were designed to amplify regions of the different Y-linked scaffolds followed by polymerase chain reaction (PCR) using either male or female genomic DNA as template. The primers were designed using the 10x-only Y scaffolds since they have the highest identity. When a primer pair resulted in the amplification of the expected size band with male genomic DNA only, we concluded that its corresponding scaffold was Y-specific. Based on this criterion, 28 scaffolds were considered Y-specific (Figure 6). However, it was expected that some primers might represent homologous regions between X and Y chromosomes and thus have a product both in male and female samples, albeit at a lower level in the females. Partial homology with autosomal sequences was also expected. While this difference cannot be detected in a normal PCR, it could be observed in a quantitative RT-PCR. Therefore, lower male qRT-PCR cycle-threshold (Ct) amplification values than female should indicate that the respective primer pair corresponded to a Y-specific scaffold. Thus, seven more scaffolds were considered Y-specific using this approach. Of the remaining primer pairs nine gave no amplification, eleven gave ambiguous results and require further examination, while 30 amplified equally male and female gDNA.

### Assembly curation and annotation

The initial assembly (Illumina-PacBio), was previously submitted to the US National Center for Biotechnology Information (NCBI) nucleotide archive with GenBank accession: GCA_001188975.2. This genome was annotated using the NCBI Eukaryotic Genome Annotation Pipeline (Gnomon) yielding the *Bactrocera oleae* Annotation Release 100 (http://www.ncbi.nlm.nih.gov/genome/annotation_euk/Bactrocera_oleae/100/). This annotation contains a total of 13936 genes and pseudogenes of which 13198, 392, 346 are predicted to be protein-coding, non-coding, and pseudogenes, respectively. Further, 2,759 genes are predicted to have variants (isoforms). In total, the *B. oleae* genome was predicted to contain 19,694 transcripts of which 18702, 411, 393, 188 are mRNA (with CDSs), tRNA, lncRNAs, and miscellaneous RNAs, respectively. Whereas the mean length of the genes and transcripts is 9,597 bp and 2,259 bp, respectively, the longest gene is 497,921 bp while the longest transcript is 59,475 bp. All alignments and genes reported in this article refer to these references: genome GCA_001188975.2 and annotation NCBI Bactrocera oleae Annotation Release 100, unless otherwise stated.

## Discussion

The advent of low cost Illumina short-read sequencing has made this technology a standard *de novo* sequencing approach combined with specialized library preparation techniques, such as mate-pair libraries to provide anchor points for scaffold construction. Unfortunately, due to the short-read nature of this technology *de novo* genome assemblies for even prokaryotes are hugely fragmented. The main bottleneck is the presence of repeated sequences in most genomes which require long-reads to span the entire repeat or provide enough resolution to unambiguously connect two contigs. More recently, long-read sequencing technologies, such as SMRT (Pacific Biosciences) and protein nanopore (Oxford Nanopore Technologies) produce long reads and thus enable construction of highly contiguous assemblies. The drawback of long-read sequencers is their relatively high error rates. Further, long-read technologies require genome coverage at approximately 60X and 30X, respectively, and with their current low throughput makes them relatively more expensive with PacBio and ONT currently costing USD 100 - 200 per Gb, which is ∼5-10 times more than Illumina^41^. However, the ONT PromethION can cost USD 12 – 24 per Gb due to increased throughput^41^. In parallel, long-read sequencing library preparation protocols put a considerable demand on DNA quantity required for constructing and sequencing long-read libraries ^42^. Typically, >1000 ng of library are needed as input in these systems compared to 0.5 – 1.2 ng needed with the 10x Genomics system.

The, Linked-reads technology (10x Genomics), has emerged as an alternative approach to obtain long-range sequence information and even construct haplotype specific assemblies while leveraging the high throughput, low cost, and accuracy of short-read sequencing. Recent *de novo* genome assembly projects have opted to combine different technologies^43–45^ highlighting the benefit of hybrid genome assembly. Particularly important is the combination of highly accurate short-read SGS technologies with long-read TGS technologies. The 10x Genomics platform presents an interesting approach of using short-read to gain long range genomic information. Here, we used the highly accurate reads from the 10x Genomics workflow to correct assemblies generated by long-read technologies, and then used the corrected assemblies to scaffold and close gaps in the 10x assembly.

The olive fruit fly (*Bactrocera oleae*) is a very important agricultural pest that causes huge economic losses to the olives industry estimated at USD 800 million per year. As part of the i5k project, we aimed to sequence the genome of the olive fly. Our initial efforts resulted in the currently available genome assembly (GenBank assembly accession: GCA_001188975.2) and annotation (*Bactrocera oleae* Annotation Release 100). However, because this genome was assembled primarily with Illumina short-reads and scaffolded with Illumina mate-pair reads and PacBio reads the contiguity was low (N50 of 0.14 Mb). In this work we sequenced the genome using linked-reads and ONT technologies. The ONT technology was particularly chosen because it has the longest read lengths and shows good ability to sequence DNA across a range of sizes yielding reads with mixed insert sizes, a feature that improves assembly quality^46^.

The linked-reads assembly was by far the most contiguous giving a scaffold N50 of 2.16 Mb compared to the ONT assembly which had 0.26 Mb. However, the 10x-only assembly had 3544 N’s per 100 kb whereas the ONT-only assembly has no gaps. Breaking the 10x-only scaffolds into contigs resulted into a contig N50 of 0.1 Mb (Supplementary Figure S6). Therefore, the genome derived from ONT reads was more contiguous than the linked-reads assembly on a contig level. Combination of ONT and PacBio reads created a more contiguous assembly. It is however, likely that just doubling our coverage for ONT reads could have had a similar effect. We then evaluated the effect of scaffolding the linked-reads assembly using Illumina mate-pairs, PacBio, and/ONT. Interestingly, scaffolding of the 10x-only assembly using ONT reads yields similar results to scaffolding the 10x-only assembly with all the platforms (contig N50 of 4.59 vs 4.89, respectively). This demonstrated the utility of the ONT platform. We therefore, find that genome assembly using the linked-reads technology followed by ONT yields highly contiguous assemblies.

Correction of errors in assemblies generated from TGS technologies is critical to obtaining a high-quality assembly. Several approaches exist but here we used short reads to do the error correction. As shown in Figure 5, error correction enables a significant improvement in representation of conserved genes. The 10x-only also showed the highest representation of ancestral conserved genes (BUSCOs) in the 4 levels tested; Eukaryota, Arthropoda, Insecta, and Diptera, recovering over 97% of the genes in each category. Due to high contiguity and low error rate in the 10x-only assembly we opted to use this assembly for further annotation.

Previous studies have shown that 50% of the human genome is composed of repetitive sequences^47^ while 29% of the *D. melanogaster* genome is composed of repetitive sequences^48^. Using RepeatMasker, we found that 12% of the olive fly genome is composed of repeats while the total length of repeats identified using RepeatScout was 34%. Interestingly though, DNA Transposons are the most prevalent repeat sequences followed by an unclassified class unlike in *D. melanogaster* where Long Terminal Repeats are the most prevalent followed by Long Interspersed Nuclear Elements (LINEs).

The Y chromosome remains as one of the most challenging chromosomes to assemble due to the highly repetitive nature of its sequence. Using the chromosome quotient method, we were able to identify putative Y chromosome scaffolds. The assembly generated by combining all technologies, 10x-all produced the most contiguous Y chromosome assembly. The largest scaffold was 318 kb and N50 of 60 kb. However, the assembly generated from long reads (ONT-PacBio) was the longest (4.72 Mb), containing no gaps, and had only 125 contigs compared to the 599 scaffolds for the 10x-all Y chromosome assembly. This highlights the utility of long-read technologies in attempting to assemble highly repetitive regions. Indeed, Miten et al used Oxford Nanopore technology to assemble for the first time an array repeat stretching ∼300 kb and spanning a human centromere on the Y chromosome^49^.

Developing a high-quality reference grade olive fly assembly has been a critical step in progressing olive fly research. Despite its economic importance, olive fly’s molecular research was lagging due to, among others, difficulty in rearing and absence of classical genetic tool. The current genome assembly offers a great leap forward in olive fly genomics research, setting a paradigm for other medium sized animal genomes. Of particular interest is the assembly of the Y chromosome where the male determining factor, M, resides. On one hand, understanding the structure and organization of olive fly’s Y chromosome will shed light on its function and evolution. On the other, the identification of the male determining factor could play a critical role in sterile insect technology (SIT) where genetically fit sterile males could be more easily and reliably produced and used to control spread of olive flies. Previous attempts at sterile insect technique and biological control failed and thus the current control measures largely rely on the use of pesticides. Providing a high-quality genome assembly will tremendously improve our understanding of the biology of this organisms and hopefully open avenues for biological control. Genome curation and annotation will be the subject of a future publication.

