## Supplemental material for "*De novo* genome assembly of the olive fruit fly (*Bactrocera oleae*) developed through a combination of linked-reads and long-read technologies"

Tabe S1: Summary of library used in the our initial assembly generated from Illumina and PacBio sequencing

Library Table

| Library ID | Sample | Library Type | Shear Size | Actual Size | Sequcing type | Reads sequenced | Bases sequenced | Coverage |
|---|---|---|---|---|---|---|---|---|
| 1 | Male gDNA | Truseq DNA (PCR) | - | - | PE100 | 114,837,736 | 11,483,373,600 | 36X |
| 2 | Female gDNA | Truseq DNA (PCR) | - | - | PE100 | 197,243,244 | 19,724,324,400 | 61X |
| 3 | Male gDNA | Nextera MP | 6.4 kb | 4 kb | PE100 | 33,643,072 | 3,364,307,200 | 10X |
| 4 | Male gDNA | Nextera MP | 9.1kb | 6.2 kb | PE100 | 26,505,296 | 2,650,529,600 | 8X |
| 5 | Male gDNA | Nextera MP | 9.8kb | 6.7 kb | PE100 | 275,608,812 | 27,560,881,200 | 86X |
| 6 | Male gDNA | PacBio | 17kb | 6 kb | P4-C6 Chemistry | 1,084,895 | 6,458,961,607 | 20X |

| Library | kmer | Total bases | Contigs | GN50 Count | GN50 Length | Largest | Best |
|---|---|---|---|---|---|---|---|
| mix | k31 | 391484560 | 146562 | 6188 | 10639 | 390431 | |
| mix | k35 | 402562855 | 140115 | 5568 | 11933 | 456670 | |
| mix | k37 | 407245803 | 137258 | 5381 | 12391 | 433281 | |
| mix | k39 | 410728391 | 135584 | 5222 | 12930 | 397852 | |
| mix | k41 | 413598928 | 134199 | 5068 | 13362 | 509383 | chosen |
| mix | k43 | 416895565 | 132926 | 4926 | 13826 | 397861 | |
| mix | k45 | 419301687 | 131769 | 4833 | 14259 | 397861 | |
| mix | k51 | 417279050 | 114295 | 4563 | 15232 | 397882 | |
| mix | k61 | 408251794 | 109474 | 4497 | 16419 | 412306 | |
| female | k31 | 380383328 | 140667 | 5772 | 11148 | 299971 | |
| female | k33 | 385548458 | 137517 | 5436 | 11910 | 309491 | |
| female | k35 | 389097923 | 135187 | 5233 | 12435 | 309490 | |
| female | k37 | 392910761 | 133447 | 4990 | 13104 | 397852 | |
| female | k39 | 395280236 | 132492 | 4865 | 13488 | 421774 | |
| female | k41 | 397620313 | 131381 | 4719 | 14041 | 433284 | chosen |
| female | k61 | 360009040 | 139790 | 7942 | 8986 | 222154 | |
| male | k31 | 365812436 | 177101 | 8673 | 8569 | 209874 | |
| male | k33 | 369580179 | 177545 | 8463 | 8859 | 232360 | |
| male | k35 | 371134473 | 179668 | 8460 | 8915 | 240645 | |
| male | k37 | 372037488 | 183157 | 8817 | 8550 | 279665 | chosen |
| male | k39 | 372362904 | 188288 | 9303 | 8154 | 231995 | |
| male | k41 | 371294702 | 194310 | 10092 | 7506 | 192760 | |
| male | k51 | 306771946 | 194321 | 26273 | 2573 | 74978 | |

| Tool | Version | Link |
|---|---|---|
| RAY | 2.2.0-k96 | http://denovoassembler.sourceforge.net/ |
| SSPACE | 2.0_Basic | http://www.baseclear.com/landingpages/basetools-a-wide-range-of-bioinformatics-solutions/sspacev12/ |
| NextClip | v0.8 | https://www.ncbi.nlm.nih.gov/pubmed/24297520 |
| bwa-mem | 0.6.2 | http://bio-bwa.sourceforge.net/ |
| Trimmomatic | 0.25 | http://www.usadellab.org/cms/index.php?page=trimmomatic |
| kmergenie | x | x |

Table S2: Quast results for 3 assemblies

| | **Assembly** | | |
|---|---|---|---|
| | **illumina-PacBio** | **10x-only** | **ONT-only** |
| # scaffolds/contigs | 36198 | 34612 | 2473 |
| Largest contig | 5074932 | 12011473 | 4384956 |
| Total length | 471780370 | 427015443 | 396506582 |
| GC (%) | 34.48 | 34.69 | 35.18 |
| N50 | 139566 | 2158770 | 267480 |
| L50 | 474 | 44 | 408 |
| # N's per 100 kbp | 10853.91 | 3543.94 | 0 |

Table S3: Tissues used in the transcriptome sequencing and assembly. The number of reads generated, and the reads paired are shown

| Sample | Raw Paired Reads | Surviving Paired Reads | % |
|---|---|---|---|
| **Bo_adult** | 34849995 | 32844054 | 94 |
| **Bo_pupa** | 40630107 | 38409178 | 95 |
| **Bo_Thorx_Mix** | 54414205 | 51572554 | 95 |
| **Bo_egg** | 37651087 | 35142389 | 93 |
| **Bo_larva** | 38678071 | 36544941 | 94 |
| **Bo_Npin_Fem** | 41300500 | 39431112 | 95 |
| **SexOrgans_male** | 33007668 | 30981652 | 94 |
| **Bo_Ypin_Fem** | 45137730 | 42459646 | 94 |
| **Bo_Cpin_Fem** | 39353529 | 36854295 | 94 |
| **Testes_after** | 34278769 | 31971074 | 93 |
| **Bo_Legs_ovip_Female** | 38111237 | 35695711 | 94 |
| **Bo_Heads_Ovip_Female** | 37064417 | 35037660 | 95 |
| **Bo_ovipositor** | 29211750 | 27366916 | 94 |
| **Bo_Heads_Female** | 36884461 | 34599100 | 94 |
| **Bo_Heads_Male** | 53245375 | 50049221 | 94 |

Table S4: Summary of the Trinity de novo transcriptome generated from sequencing all the tissues in Table S3.

| Transcript Length Distribution | |
|---|---|
| Description | Value |
| Nb. Transcripts | 133003 |
| Nb. Components | 85261 |
| Total Transcripts Length (bp) | 146616389 |
| Min. Transcript Length (bp) | 201 |
| Median Transcript Length (bp) | 503 |
| Mean Transcript Length (bp) | 1102 |
| Max. Transcript Length (bp) | 27500 |
| N50 (bp) | 2266 |

Table S5: Assessment of the completeness of the Trinity de novo transcriptome assembly. The assessment was carried out using gVolante[1] selecting for Arthropoda BUSCOs.

| | |
|---|---|
| Total number of core genes queried | 1066 |
| Number of core genes detected | |
| Complete | 1054 (98.87%) |
| Complete + Partial | 1062 (99.62%) |
| Number of missing core genes | 4 (0.38%) |
| Average number of orthologs per core genes | 1.65 |
| % of detected core genes that have more than 1 ortholog | 44.02 |
| Scores in BUSCO format | C:98.8%[S:55.3%,D:43.5%],F:0.8%,M:0.4% |

Table S6: Quast results for 3 Y chromosome assemblies; an assembly generated from the 10x Genomics linked-reads technology (Y-10x-only), a hybrid assembly generated only from Oxford Nanopore technology and PacBio (Y-ONT-PacBio), and an assembly generated from the 10x Genomics linked-reads technology scaffolded with all other technologies (Y-10x-all).

| | **Assembly** | | |
|---|---|---|---|
| | **Y-10x-only** | **Y-10x-all** | **Y-ONT-PacBio** |
| # contigs/scaffolds | 2138 | 873 | 125 |
| Largest contig (bp) | 177945 | 318374 | 204928 |
| Total length | 3650627 | 3802767 | 4721026 |
| GC (%) | 32.14 | 33.82 | 46.24 |
| N50 | 3337 | 60003 | 43122 |
| L50 | 138 | 13 | 38 |
| # N's per 100 kbp | 55238.98 | 75614.73 | 0 |

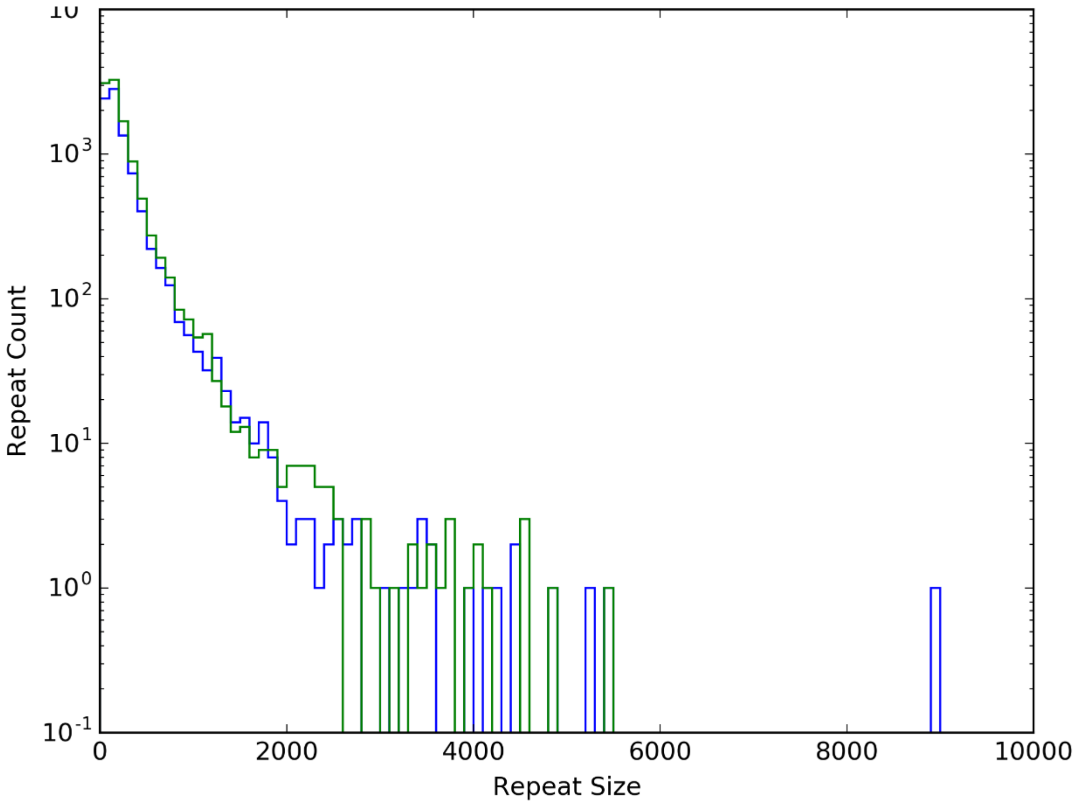


Figure S1: Repeat content of 2 genomes as identified by RepeatScout. The assembly generated 10x Genomics linked-reads technology alone (10x-only, blue) and the 10x-only assembly gap-closed using PacBio reads (green) are shown.

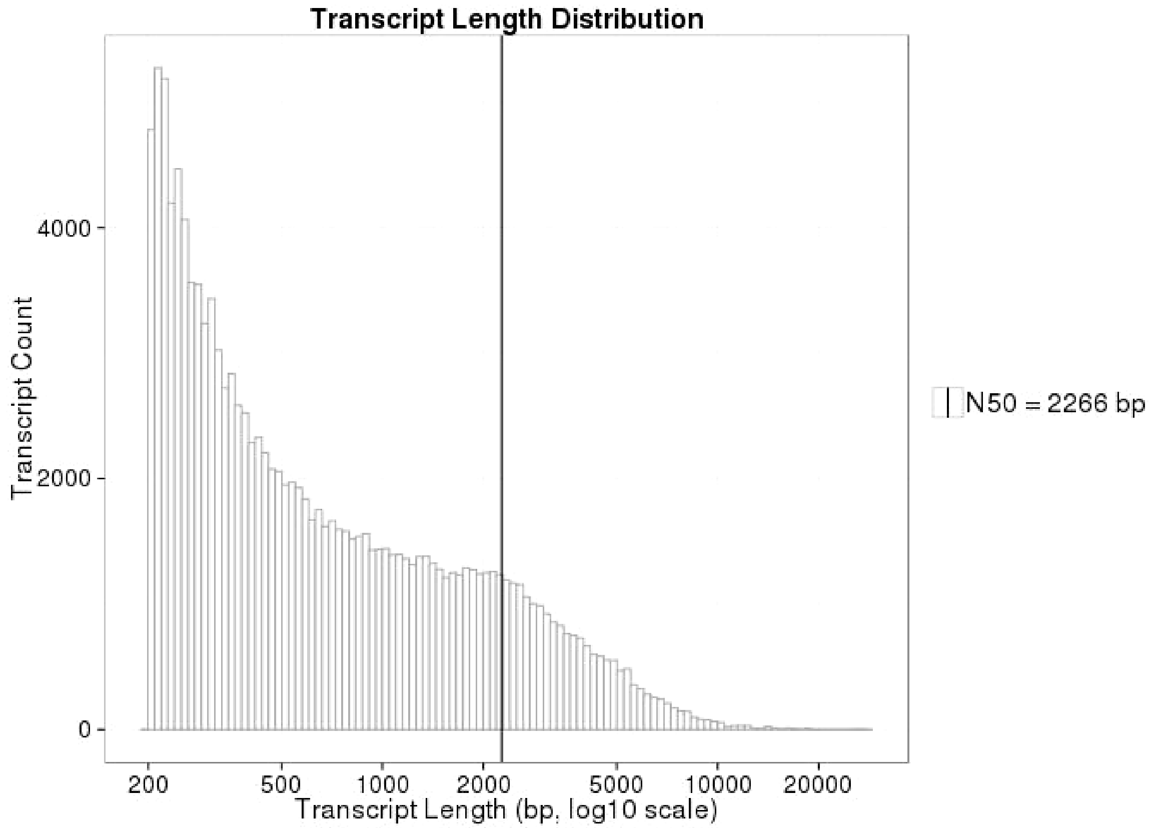


Figure S2: Histogram of transcriptome read lengths. RNA was extracted from tissues shown in Table S3 and assembled using Trinity. The histogram shows the read lengths of those transcripts.

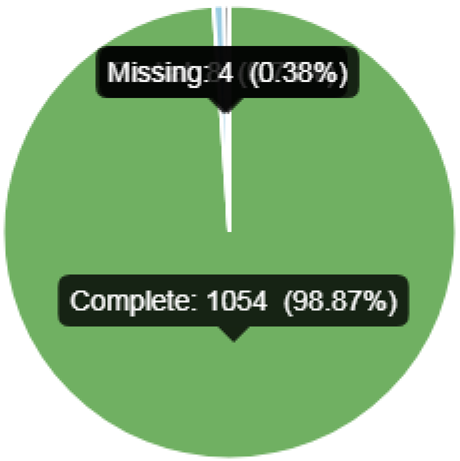


Figure S3: Assessment of the completeness of the Trinity de novo transcriptome assembly. The assessment was carried out using gVolante[1] selecting for Arthropoda BUSCOs.

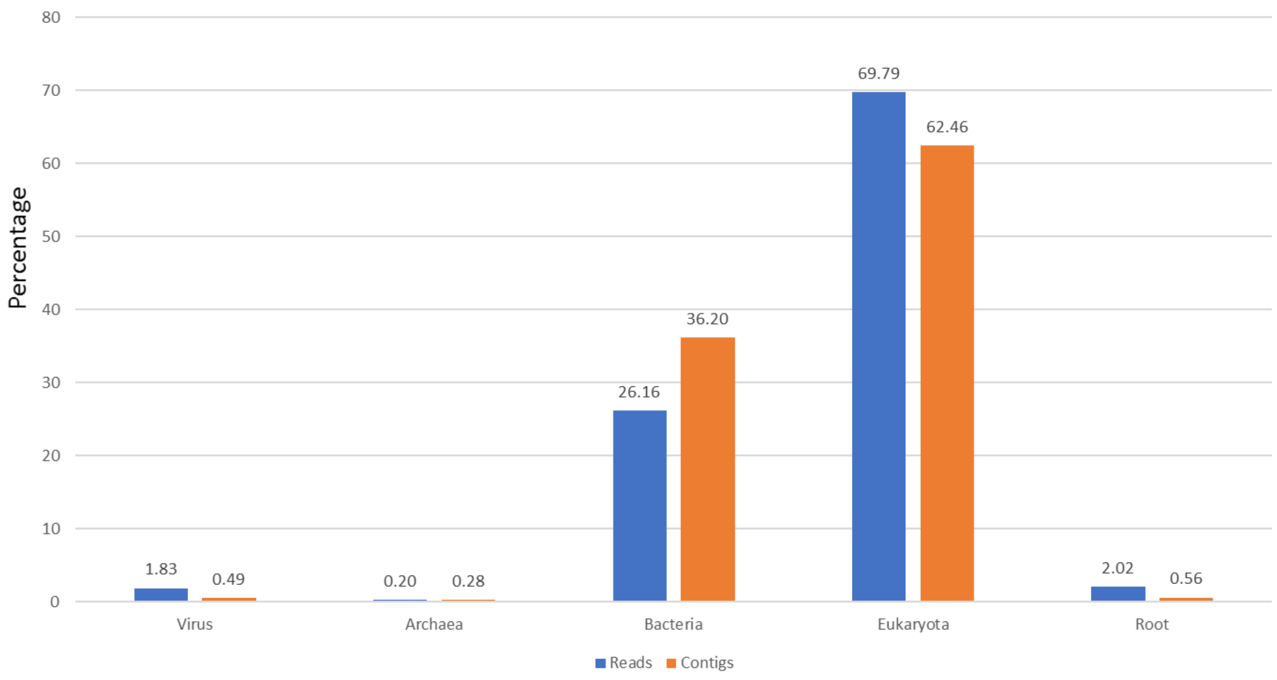


Figure S4: Bar graph showing percentage of contamination in reads (blue) and scaffolds (orange) from the 10x-only assembly. Reads and scaffolds were analysed using Kraken[2]

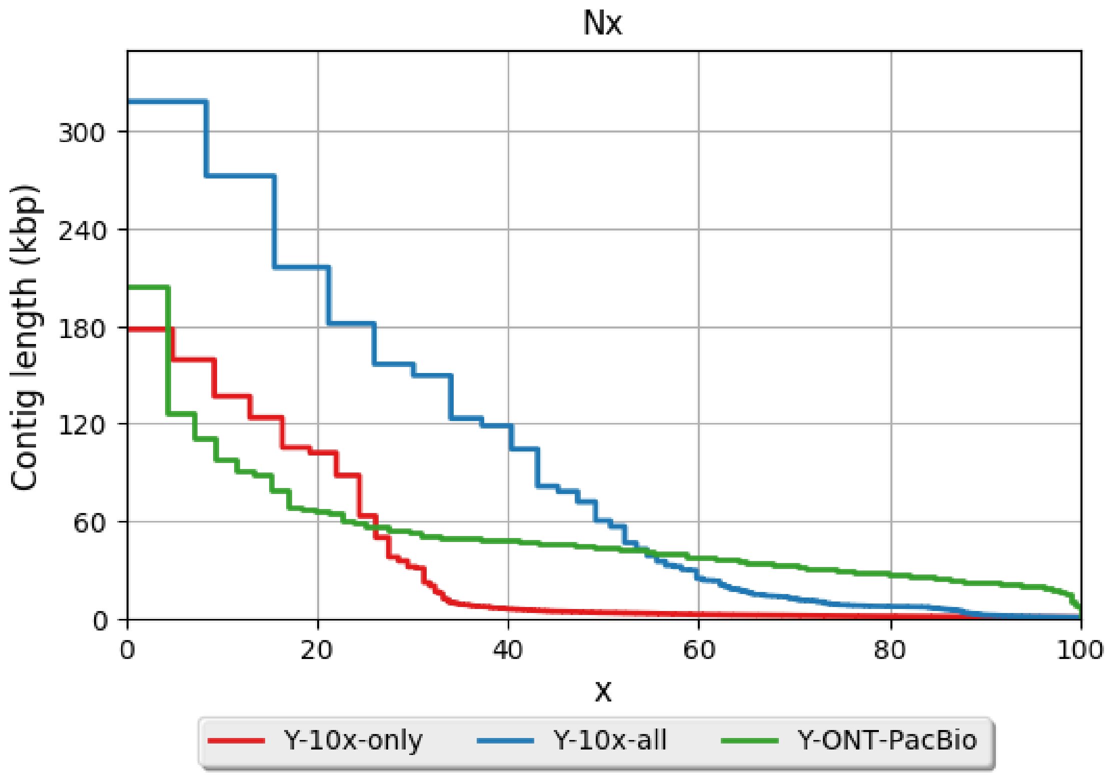


Figure S5: Contiguity plot generated using Quast[3] showing results for 3 Y chromosome assemblies; an assembly generated from the 10x Genomics linked-reads technology (Y-10x-only), a hybrid assembly generated only from Oxford Nanopore technology and PacBio (Y-ONT-PacBio), and an assembly generated from the 10x Genomics linked-reads technology scaffolded with all other technologies (Y-10x-all). When all contigs from an assembly are ordered from largest to smallest, x is described as the value at which x% of the genome is contained in contigs at and above this contig length. The y-axis shows the contig/scaffold length at each of the Nx values. The N50 is the contig/scaffold length at which 50% of the genome is contained in scaffolds bigger as big as that contig/scaffold.

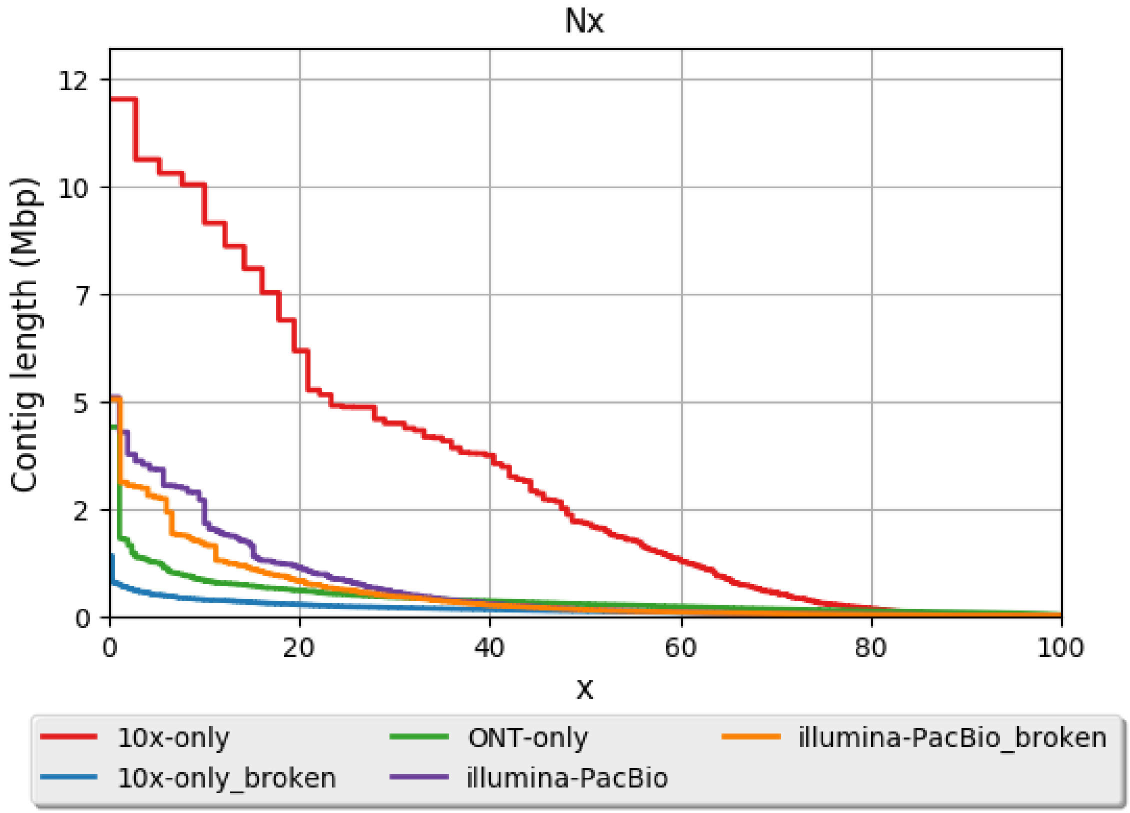


Figure S6: Contiguity plot generated using Quast[3] showing results for different assemblies. The assemblies ending with broken are derived from breaking the scaffolds where gaps exist to create contigs. When all contigs from an assembly are ordered from largest to smallest, x is described as the value at which x% of the genome is contained in contigs at and above this contig length. The y-axis shows the contig/scaffold length at each of the Nx values. The N50 is the contig/scaffold length at which 50% of the genome is contained in scaffolds bigger as big as that contig/scaffold.
